# Influenza A defective viral genome production is altered by metabolites, metabolic signaling molecules, and cyanobacteria extracts

**DOI:** 10.1101/2024.07.04.602134

**Authors:** Ilechukwu Agu, Ivy R. José, Samuel L. Díaz-Muñoz

**Affiliations:** Department of Microbiology and Molecular Genetics University of California, Davis, One Shields Ave Davis CA 95616; Genome Center, University of California, Davis One Shields Ave, Davis CA 95616

## Abstract

RNA virus infections are composed of a diverse mix of viral genomes that arise from low fidelity in replication within cells. The interactions between “defective” and full-length viral genomes have been shown to shape pathogenesis, leading to intense research into employing these to develop novel antivirals. In particular, Influenza A defective viral genomes (DVGs) have been associated with milder clinical outcomes. Yet, the full potential of DVGs as broad-spectrum antivirals remains untapped due to the unknown mechanisms of their *de novo* production. Much of the research into the factors affecting defective viral genome production has focused on the virus, while the role of the host has been neglected. We recently showed that altering host cell metabolism away from pro-growth pathways using alpelisib increased the production of Influenza A defective viral genomes. To uncover other drugs that could induce infections to create more DVGs, we subjected active influenza infections of the two circulating human subtypes (A/H1N1 & A/H3N2) to a screen of metabolites, metabolic signaling molecules, and cyanobacteria-derived biologics, after which we quantified the defective viral genomes (specifically deletion-containing viral genomes, DelVGs) and total viral genomes using third generation long-read sequencing. Here we show that metabolites and signaling molecules of host cell central carbon metabolism can significantly alter DelVG production early in Influenza A infection. Adenosine, emerged as a potent inducer of defective viral genomes, significantly amplifying DelVG production across both subtypes. Insulin had similar effects, albeit subtype-specific, predominantly enhancing polymerase segment DVGs in TX12 infections. Tricarboxylic Acid (TCA) cycle inhibitors 4-octyl itaconate and UK5099, along with the purine analog favipiravir, increased total viral genome production across subtypes. Cyanobacterial extracts primarily affected DVG and total viral genome production in TX12, with a specific, almost complete shutdown of influenza antigenic segments. These results underscore the influence of host metabolic pathways on DVG production and suggest new avenues for antiviral intervention, including PI3K-AKT and Ras-MAPK signaling pathways, TCA cycle metabolism, purine-pyrimidine metabolism, polymerase inhibition, and cyanotherapeutic approaches. More broadly, our findings suggest that the social interactions observed between defective and full-length viral genomes, depend not only on the viral actors, but can be altered by the stage provided by the host. Our study advances our fundamental understanding of DVG production mechanisms and highlights the potential of targeting host metabolism to develop broad-spectrum influenza therapeutics.

## Introduction

RNA virus infections are not homogenous, instead consisting of a diverse mix of virions with substantial genetic variation. Defective viral particles, named for the defective viral genomes they harbor, are an exceedingly common part of this mix, often outnumbering “standard” virions and genomes. Aside from their abundance, defective viral particles have long been of interest because they require complementation from “standard” viral genomes, yet in many cases they potently interfere with the replication of these very standard genomes. These interactions between virions lead to social dynamics (Díaz-Muñoz et al. 2017) that alter the composition of infections. Recently, defective viral particles have been identified in clinical settings and associated with milder infection outcomes (Vasilijevic, 2017). Intense research efforts have focused on understanding the mechanisms of defective virus emergence and on developing novel, broad-spectrum antivirals.

Influenza A virus remains a substantial public health threat despite yearly vaccination efforts (Rolfes, 2019), which are thwarted by low vaccination rates (Chen, 2022) and high mutability of Influenza A (Pauly, 2017). In the face of antivirals that have not been effective (Hanula, 2024) or that have been rendered ineffective through antiviral resistance (Hurt, 2012; Dong, 2015), defective viral genomes as therapeutics have emerged as a promising alternative. Defective viral genomes in influenza are **Del**etion-containing **V**iral **G**enome (hereafter DelVGs after Alnaji et al. 2019) segments which are packaged into defective interfering particles (Henle, 1943; Von Magnus, 1954; Davis, 1980; Nayak, 1982; Saira, 2013) that hinder productive pathogenesis if they accumulate naturally (Dimmock, 2014; Manzoni, 2018; Vignuzi, 2019; Wu, 2022) or are exogenously introduced (Smith, 2016; Meng, 2017; Wasik, 2018; Zhao, 2018; Bdier, 2019; Yamagata, 2019; Tapia, 2019; Harding, 2019). However, full weaponization of DelVGs against the flu in a broad-spectrum manner remains unrealized because their *de novo* mechanism of production is unknown, and few factors associated with their production are known (Von Magnus, 1954; Bangham, 1990; Perez-Cidoncha, 2014; Vasilijevic, 2017; Agu et al., 2024). Thus, progress in uncovering novel effectors of DelVG *in situ* production has been slow. There is a critical need to discover novel effectors of DVG production so that the shared characteristics of these inducers may inform the molecular-level events proximal to DVG production.

Research into Influenza A defective interfering particles and their DelVGs has primarily focused on viral factors affecting DelVG production, such as multiplicity of infection, polymerase gene mutations, and each strain’s propensity to generate DelVGs (Alnaji, 2019). At a molecular and cellular level, substantial research has focused on polymerase synthesis (Nayak, 1982; Winters, 1981; Alnaji, 2020) and differential packaging (Brooke, 2014; Alnaji, 2021; Meng, 2017; Odagiri, 1997) of defective viral genome segments. Further, there has been considerable effort into the effects that DelVGs have on the host (De, 1980; Tapia, 2013; Frensing, 2014; Vasilijevic, 2017; Wang, 2023). Surprisingly, the potential role of the *host* in influencing DelVG composition has been largely overlooked (Ginsberg, 1954; Choppin, 1969; Choppin, 1970; De, 1980). Host cell metabolism and metabolic signaling, can affect progeny virus yield and the severity of infection (Ackermann, 1951; Kilbourne, 1959; Blough, 1963; Hale, 2006; Li, 2008; Kuss-Duerkop 2017; Ren, 2021), leading us to investigate the impact of host metabolic signaling on the production of Influenza A defective particles and DelVGs. We recently showed that Alpelisib—a highly selective inhibitor of mammalian phosphoinositide-3 kinase (PI3K) and its downstream metabolic signaling (Furet, 2013; Fritsch, 2014)— increased DelVG production during Influenza A infection (Agu et al., 2024). On the basis of this association between host metabolism and DelVG production, and the extensively characterized crosstalk between host metabolism and flu pathogenesis, we hypothesized that other metabolites, metabolic signaling molecules, or other compounds capable of causing metabolic perturbations could increase *de novo* DelVG production in cells.

We set out to test this hypothesis by using third-generation, long-read sequencing to monitor how DelVG production was affected early in infections with A/California/07/2009 (H1N1) and A/Texas/50/2012 (H3N2) strains (CA09 and TX12 hereafter). The metabolic pathways of interest we selected, and their associated drugs, include PI3K-AKT pathway signaling (insulin, alpelisib, MK2206), tricarboxylic acid (TCA) cycle (UK5099, 4-octyl itaconate), and purine-pyrimidine metabolism (adenosine, uridine). We included favipiravir (T-705) in our screen to uncover if its role as a mutagen and elongation terminator of RNA polymerization could impact DVG production; it is a purine base analog that is converted into nucleoside and nucleotide-phosphate forms within cells (Shiraki, 2020). We also included cyanobacteria extracts from *Leptolyngbya*, *Nostoc*, *Oscillatoria, Synechococcus*, and *Tolypothrix* to uncover if the known effects of some cyanobacteria extracts on flu pathogenesis (Singh, 2017; Silva, 2018; Mazur-Marzec, 2021) extend to altered DelVG production.

We show that adenosine is a potent amplifier of DelVG production across subtypes, with the largest and most consistent increases of any compound we tested. Meanwhile, the increased DelVG production associated with insulin was more restricted to the polymerase segments and more pronounced for TX12 than CA09. Additionally, we found that TCA cycle flux inhibitors 4-octyl itaconate (4-OI) and UK5099 are potent amplifiers of total viral genome production across subtypes, and that the purine analog favipiravir also increased total viral genomes across subtypes. Finally, we discovered that cyanobacterial extracts affected defective and total viral genomes primarily in TX12. These results strengthen our previously established association between host metabolism and DelVG production (Agu et al., 2024), further cementing host involvement in the emergence of DelVGs. These results also reveal viral polymerase inhibition and cyanotherapeutic intervention as novel means of altering DelVG production, and steering the trajectory of Influenza A infection. Altogether, our findings mark a step forward in understanding the mechanism of *de novo* DVG production by identifying related environmental effectors. The shared characteristics of these effectors may shed light on the molecular events leading to DVG production and guide highly targeted future research into these mechanisms.

## Methods

### Cells and Viruses

We obtained MDCK-London cells from the Unites States Centers for Disease Control and Prevention (CDC) Influenza Reagent Resource (IRR). We maintained cells in minimum essential media (MEM) plus 5% fetal bovine serum (FBS). Egg-passaged wildtype A/California/07/2009 (H1N1) and A/Texas/50/2012 (H3N2) were a gift from the lab of Dr. Ted Ross. These initial stocks were double plaque purified in MDCK cells (ATCC/BEI) and propagated thereafter at low infection multiplicity (0.001) in MDCK cells (ATCC/BEI).

### Cyanobacteria Culture and Lysis

We cultured *Nostoc punctiforme* (ATCC 29133) in fourfold diluted Allen & Arnon liquid media (AA/4) minus ammonium or other nitrogen source (Allen, 1955); so as to induce heterocyst formation. We cultured *Leptolyngbya boryana* PCC 6306, *Oscillatoria* sp. (ATCC 27930), *Synechococcus elongatus* PCC 7942, and *Tolypothrix* sp. PCC 7601 in BG11 media. All cyanobacteria cultures were incubated at room temperature on shaker platforms (75-100 rpm), and illuminated with three overhead 20-Watt cool-white fluorescent bulbs.

We transferred 6.5 – 7.5 g of each cyanobacteria specimen along with growth media into a sealed centrifuge tube and incubated them for 24 hr at room temperature, in the dark, without shaking. Next, we washed cyanobacteria specimens once in PBS, transferred ∼1 g into 1 mL dH2O, and then froze specimens at -80 °C for 48 hr. After the freezing period, we thawed specimens at room temperature for 24 hr and filter-sterilized the lysate with a 0.22 micron syringe filter. We confirmed lysis and cell death in four of the five specimens; firstly via strain-specific coloration of the lysate (*Leptolyngbya* – cobalt blue, *Nostoc* – purple, *Oscillatoria –* clear/no lysis, *Synechococcus –* pale yellow-green, *Tolypothrix* – indigo), and secondly under the light microscope. Despite the *Oscillatoria* filaments remaining intact, we proceeded with its conditioned freeze-thaw vehicle media (dH2O).

### Drug Screening Assay

We seeded MDCK-London cells overnight in MEM plus 5% FBS media for 21 hr into plastic-bottom 96- well tissue culture plates. We then serum-weaned the fully confluent monolayers for 3 hr in MEM plus 5% bovine serum albumin (BSA). At the start of serum weaning, we spiked a 10 uL pre-treatment of vehicle control (DMSO or dH2O), cyanobacteria freeze-thaw extract (*Leptolyngbya*, *Nostoc*, *Synechococcus*, *Tolypothrix*), cyanobacteria-conditioned freeze-thaw vehicle medium (*Oscillatoria*), or 21X of small molecule drug (insulin, alpelisib, MK2206, adenosine, uridine, UK5099, 4-octyl itaconate, or favipiravir) directly into 200 uL of the serum weaning supernatant to reach a 1X concentration (1 µM). At the end of serum weaning, we exchanged serum weaning media with MEM plus 2% BSA and 1% Anti-Anti (Virus Infection Media; VIM). We then mock-infected monolayers with VIM, or we virus-infected monolayers at a multiplicity (MOI) of 1; no trypsin was used. As part of the inoculation regimen, we spiked 1.9 uL of vehicle control, cyanobacteria extract, or 21X of signaling molecule into 40 uL of the inoculum supernatant to reach a 1X concentration (1 µM), in order to sustain each treatment’s effects for the duration of virus-monolayer adsorption. After the 1 hr adsorption incubation, we aspirated inocula, washed monolayers and topped them with 200 uL VIM, and we spiked 10 uL of vehicle control, cyanobacteria extract, or 21X of signaling molecule directly into VIM supernatant to reach a 1X concentration (1 µM). After 17 hr post infection, we harvested and titrated supernatants to determine the total number of viral genomes, as well as the relative abundances of DVGs versus full-length viral genomes. We ran three biological replicates of the experiment on different days.

### Viral genome sequencing by Nanopore long-read sequencing

We conducted our influenza A viral genome sequencing protocol that sequences and quantifies full-length and deletion-containing viral genome segments, which we have described (Agu et al., 2024). Briefly, we began by isolating viral genomic RNA from 100µL of treatment group supernatants (Zymo Research, Quick-DNA/RNA Viral MagBead kit R2140). Next, we used a 2-cycle RT-PCR reaction (Invitrogen SuperScript™ III One-Step RT-PCR System with Platinum™ Taq DNA Polymerase kit 12574026) to reverse transcribe viral genomic RNA into the first cDNA strand (1st PCR cycle), and then synthesize the second cDNA strand (2nd PCR cycle). The RT reaction to produce the first cDNA strand was primed with a 45bp forward primer (Integrated DNA Technologies) that included a complementary sequence to the uni12 region shared by all flu genomic segments (12bp), flanked with a unique molecular identifier (UMI) sequence (12bp) and a landing pad sequence for downstream barcoding primers (21bp): fwd 5’- TTTCTGTTGGTGCTGATATTGNNNNNNNNNNNNAGCRAAAGCAGG-3’.

The PCR reaction to generate the second cDNA strand was primed with a 47bp reverse primer that included a complementary sequence to the uni13 region shared by all flu genomic segments (13bp), flanked by a UMI sequence (12bp) and the barcoding primer landing pad sequence (22bp): rev 5’- ACTTGCCTGTCGCTCTATCTTCNNNNNNNNNNNNAGTAGAAACAAGG-3’. We used AMPure XP beads (Beckman Coulter, AMPure XP A63881) with manufacturer’s instructions to remove excess primers, followed by a 17-cycle amplification PCR (Invitrogen, Platinum SuperFi Master Mix 12358-050) of the umi-tagged reads with barcoding primers (Oxford Nanopore Technologies, PCR Barcoding Expansion 1-96 kit EXP-PBC096). The low number of cycles was designed to minimize PCR duplicates. We then pooled 60 ng of barcoded amplicons from each sample, cleaned and concentrated this pooled sample (Zymo Research, Select-A-Size DNA Clean & Concentrator D4080), and prepared a sequencing library in accordance with manufacturer instructions (Oxford Nanopore Technologies, Ligation-Sequencing-Kit-V14 SQK-LSK114).

We loaded the pooled libraries into an R10 flow cell connected to a MinION MkIB device and ran a 72 hr sequencing protocol from the MinKNOW control software. Upon sequencing run termination, we used the Guppy basecaller software to barcode-demultiplex sequenced reads into their respective treatment groups.

### Classification and quantification of full-length and internal-deletion viral segments

We describe the pipeline in brief below; we have published on this methodology (Agu et al. 2024) and the code for this manuscript is posted on GitHub (https://github.com/pomoxis/drug-screen-dvg). Demultiplexed amplicon sequences underwent quality control pre-processing prior to deduplication into representative sequences, after which representative sequences were classified into subgroups for DelVGs and full-length, standard viral genomes (SVG).

*Quality Control:* Our sequencing library preparation strategy began with a 1-cycle each RT-PCR then PCR addition of 12bp-long UMI sequences to the 3’ and 5’ termini of viral genomic RNA, followed by PCR addition of sequencing barcodes to both termini:

5’-barcode—spacer—landing.pad—UMI—uni12—*locus*—uni13—UMI—landing.pad—spacer—barcode-3’ For quality control, we trimmed off barcode and barcode landing pad regions with C*utadapt*, then used *Cutadapt* once more to filter-in only amplicons with a 12bp-long UMI region. Finally, we confirmed the presence of well-formed uni primer regions in the filtered amplicons before advancing to UMI deduplication.

### UMI Deduplication

We used *UMI-Tools* to sequentially group PCR duplicates by UMI, and then collapse them into a single representative read. In the final quality control step, we trimmed the uni primer region off representative reads with *Cutadapt*. By integrating UMI-deduplication into our workflow, we’ve mitigated the impact of PCR amplification bias on sequencing depths. Consequently, our DelVG and SVG count data represent a quantitative measurement of the abundance of RNA molecules (genome segments) from which the amplicons were derived.

### DelVG Characterization

UMI-deduplicated fastq files containing read sequences were processed with the Virus Recombination Mapping (*ViReMa*) software to identify recombination events per genomic read using the following parameters:

--Seed 25 --MicroInDel_Length 20 --Aligner bwa --ErrorDensity 1,25

Additionally, the *-ReadNamesEntry* switch was included in a separate ViReMa run of the same dataset in order to assign read name information to each recombination, which allowed us to collapse deletion events with the same read name into a single DelVG observation using custom Bash and AWK scripts.

--Seed 25 --MicroInDel_Length 20 --Aligner bwa --ErrorDensity 1,25 -ReadNamesEntry

### Full-length Viral Genome Characterization

To characterize SVGs, we began by using the *bwa* alignment tool to determine the properties of reads and their alignment to the reference genome; this information is captured in the bitwise FLAG field (column 2) of the output SAM file. Next, we used the *AWK* program to select only reads with proper alignment to the forward and reverse strands of the reference genome—bitwise FLAGs 0x0 (0) and 0x10 (16) respectively—and used *AWK* yet again to filter reads that were within ±100bp the length of the reference genomic segment.

## Statistics

In general, we relied on linear or linear mixed models to test significance between treatments using base R and the nmle packages, respectively. Owing to the intrinsic heterogeneity of flu infections (Russell, 2018; Wang, 2020), we included bioreplicates (three independent replicate infections conducted on different days) as a random factor in our models, unless otherwise indicated. We tested two response variables, the proportion of **Del**etion-containing **V**iral **G**enomes (DelVGs, after Alnaji et al. 2019) and the **T**otal **V**iral **G**enome count (TVG). In all cases we used per-segment data on DelVGs and TVGs, i.e. each infection had eight TVG and DelVG measurements one, per viral genome segment. We first tested for *any* statistically significant changes in TVG and DelVGs according to drug treatment. To investigate changes in DelVGs and TVGs produced by each genome segment (which have known differences in TVGs and DelVGs), we then proceeded to make models that tested each viral genome segment against the respective vehicle mock infections. When we found that strain was a significant predictor, we ran separate models to estimate coefficients and significance. We only report statistically significant results in the main text, unless otherwise noted. Code for statistics and analyses is posted on GitHub (https://github.com/pomoxis/drug-screen-dvg).

## Results

### Sequencing and measurement of influenza A viral genomes is highly consistent

We sequenced supernatants of all infections in a single MinION flow cell, which yielded 12.02M reads. After quality control for well-formed amplicons (perfect UMIs, Influenza A-specific terminal uni12/13 regions) and de-deduplicating unique molecular identifiers (UMI’s) we obtained 67,496.81 ± 56,183.60 reads per sample; note that these reads cover the entire genome segment (whether deletion-containing or not) and reflect a quantitative estimate of the vRNA molecules present in the supernatants.

We conducted mock infections using the vehicle for the drugs we screened, namely dH20 and DMSO, to serve as a control and baseline for comparison. Despite high expected variability in individual infections, especially for the production of DelVGs, our control infections with vehicle were highly repeatable in terms of the total viral of genomes (**Figure 1**) and the proportion of DVGs.

**Figure 1.**
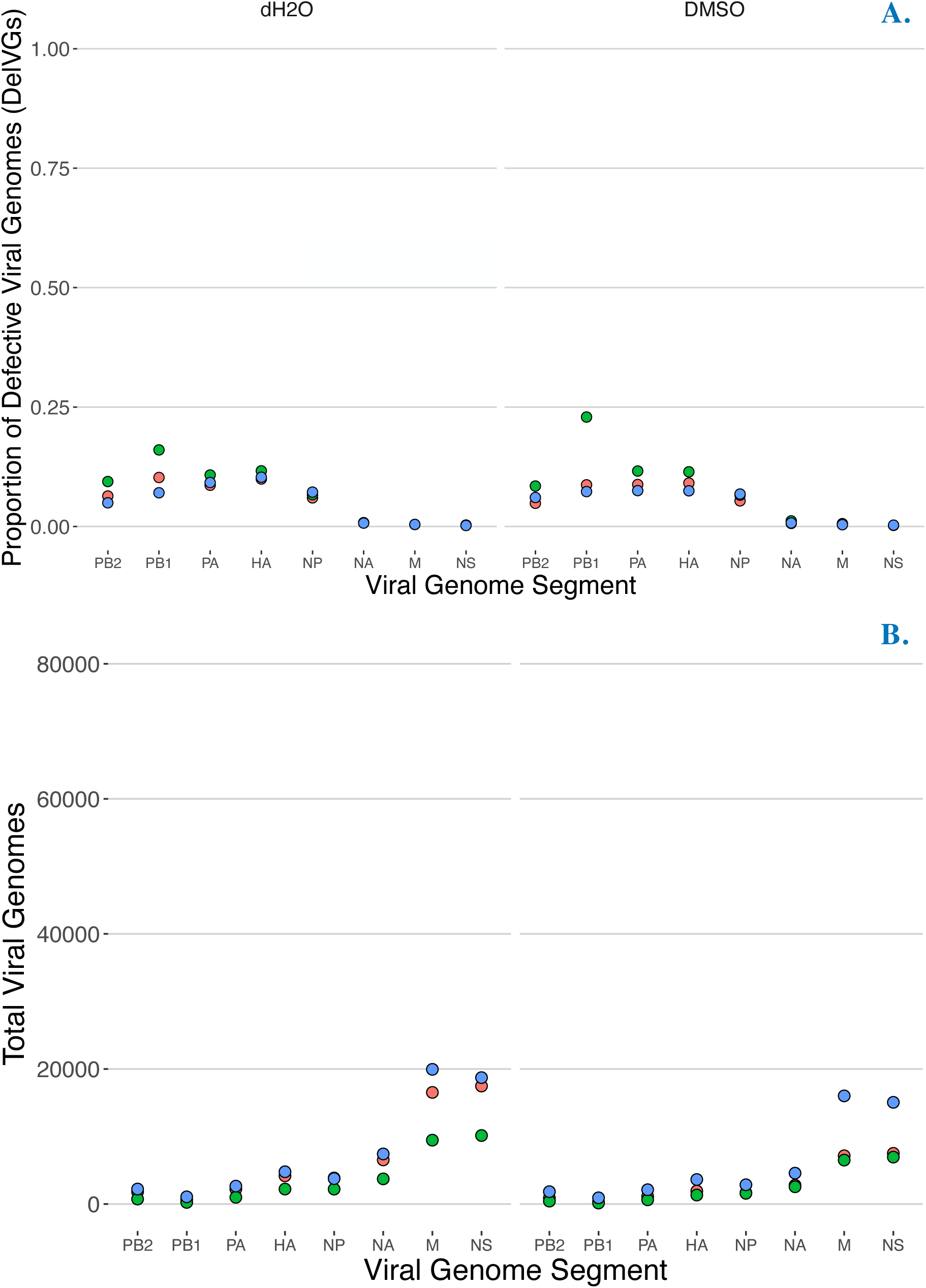
Control infections were highly repeatable. A. Production of influenza defective viral genomes (here measured as proportion of **Del**etion-containing **V**iral **G**enomes (DelVGs) in strain CA09 (H1N1pdm) after 18 hr infection (no trypsin) mock-treated with water vehicle (left panel) and DMSO vehicle (right panel). **B.** Production of influenza **T**otal **V**iral **G**enomes (TVG) in strain CA09 (H1N1pdm) after 18 hr infection (no trypsin) mock-treated with water vehicle (left panel) and DMSO vehicle (right panel). Point colors represent data from three independent infections conducted on different days (i.e. bioreplicates).

### Drugs affect defective viral genome production and total viral genome production

To determine if any of our treatments affected the proportion of DelVGs produced during an infection, we analyzed differences among treatments using ANOVA, adding bioreplicate and strain as covariates. Drug treatment was a statistically significant predictor (F = 6.777, p < 0.0001), explaining 13.12% of the variance in the proportion of DelVG among treatments. All statistically significant treatments increased the overall mean percentage of DelVGs from *on average* 8.17% to 25.61% (**Table 1**, blue shading), relative to infections with water. The largest increase was caused by Adenosine followed by Insulin, MK2206, and Uridine (in descending order, **Figure 2**; **Table 1**). We note that in this model, strain was not a significant predictor of average DelVG proportion across segments (F = 0.480, p < 0.488) and thus these average increases were significant for both TX12 and CA09 strains.

**Figure 2.**
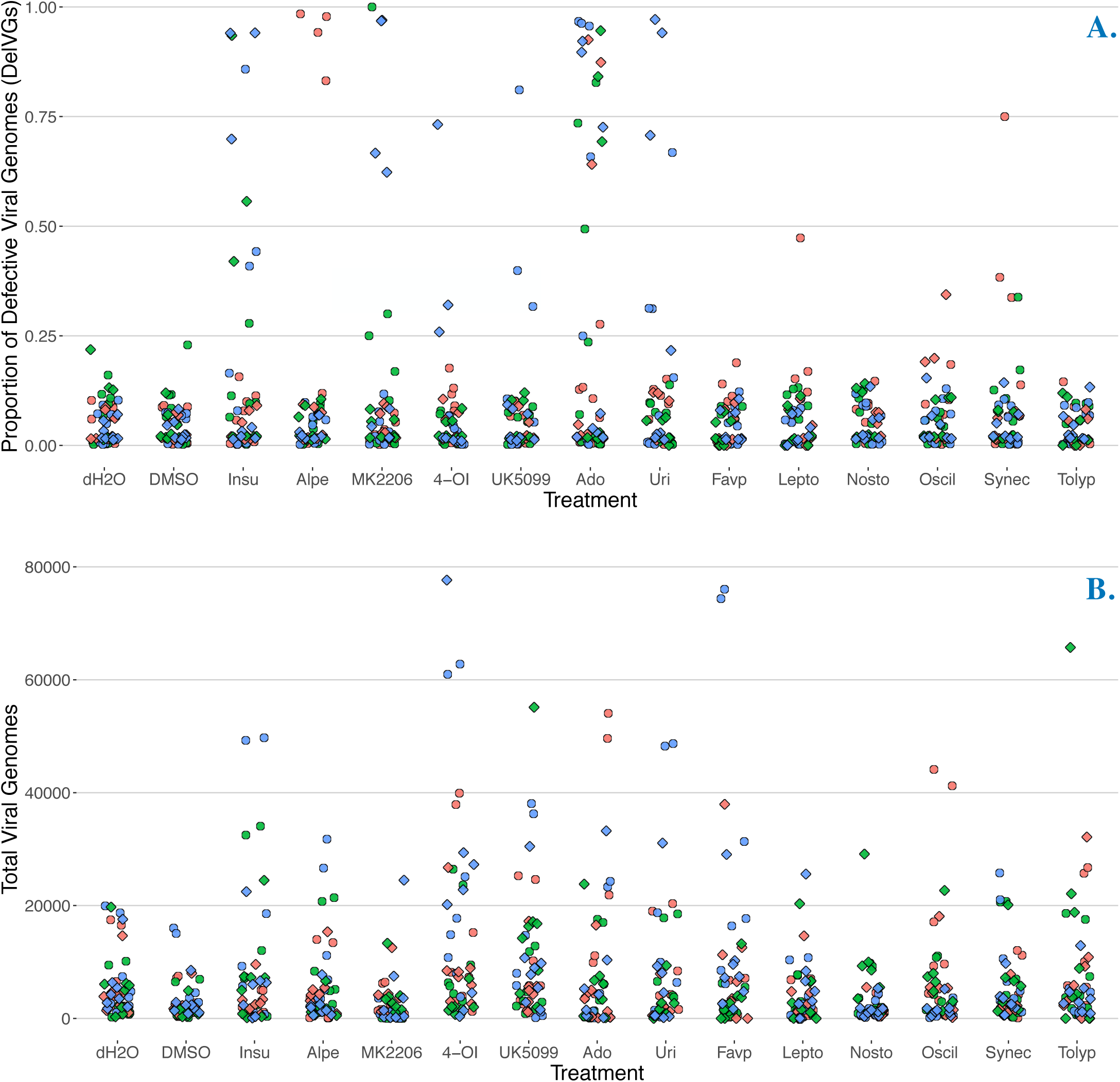
Drug treatment affects defective viral genomes and total viral genomes. A. Production of influenza defective viral genomes (here measured as proportion of **Del**etion-containing **V**iral **G**enomes (DelVGs) after 18 hr infection; each point is the proportion of DelVGs in each influenza segment (all segments included in each treatment). **B.** Production of influenza **T**otal **V**iral **G**enomes (TVG) after 18 hr infection; each point is the total number of viral genomes of each influenza segment (all segments included in each treatment). The first two treatments (left to right) are vehicle control infections. Point colors represent data from three independent infections conducted on different days (i.e. bioreplicates). The shape of the points represents the strain (circles = CA09(H1N1pdm), diamonds = TX12(H3N2)). The position of points is randomly jittered horizontally for better visibility.

**Table 1.** Drugs increase total viral genomes and deletion containing viral genomes. . Statistically significant predictors of the proportion of **Del**etion-containing **V**iral **G**enomes (DelVGs, blue shading) and **T**otal **V**iral **G**enomes (TVG) and their parameter estimates from ANOVA.

| Treatment | Measurement | Estimate | Standard Error | Pr(>\|t\|) |
| --- | --- | --- | --- | --- |
| Adenosine | Deletion-containing Viral Genomes (proportion) | 0.256059 | 0.035974 | 2.73e-12 |
| Insulin | Deletion-containing Viral Genomes (proportion) | 0.118392 | 0.035974 | 0.00105 |
| MK2206 | Deletion-containing Viral Genomes (proportion) | 0.081722 | 0.035974 | 0.02341 |
| Uridine | Deletion-containing Viral Genomes (proportion) | 0.082837 | 0.036573 | 0.02382 |
| 4-OI | Total Viral Genomes (count) | 8438.81 | 2040.61 | 3.97e-05 |
| UK5099 | Total Viral Genomes (count) | 4355.15 | 2040.61 | 0.03317 |

We similarly analyzed the number of total viral genomes produced during an infection and found that drug treatment was a statistically significant predictor (F = 4.439, p < 0.0001), explaining 8.01% of the variance in number of total viral genomes among treatments. The TCA cycle flux inhibitors 4-OI and UK5099 were statistically significant predictors of total viral genomes (**Table 1**, gray shading), increasing TVG count on average by 8,438.81 and 4,355.15 genomes, respectively, relative to infections with water (**Figure 2**; **Table 1**). Unlike the proportion of DelVGs, strain was a statistically significant predictor of total viral genomes across segments (F = 6.168, p < 0.0132). TX12 had on average -1,854.71 fewer total viral genomes than CA09.

In these two overall models, we did not control for known differences in the proportion of DelVGs in different genome segments (i.e. polymerase complex segments have a higher proportion of DelVGs). To gain insight into the segment-specific effects of each drug, we made linear mixed models for each segment and each vehicle control (DMSO or dH2O). We report the results for these models below in dedicated sections for each drug and in the supplementary information (see Results and Supplementary Material sections).

### Adenosine is a potent amplifier of DelVG production across subtypes

Adenosine had the most marked and consistent effect of any of the drugs. Adenosine increased the DelVG proportion in all three polymerase complex segments of the H1N1 and H3N2 strains in a statistically significant way (**Figure 3**, **Table S1**). The magnitude of these increases was substantial, as the average increase in polymerase complex DelVGs ranged from 35.00% to 80.61% compared to the dH2O vehicle. Adenosine also affected viral genome segments outside the polymerase complex, yielding a statistically significant increase of HA of 33.71% in CA09 infections and a small but statistically significant increase of 0.99% in NP DelVGs in TX12 infections.

**Figure 3.**
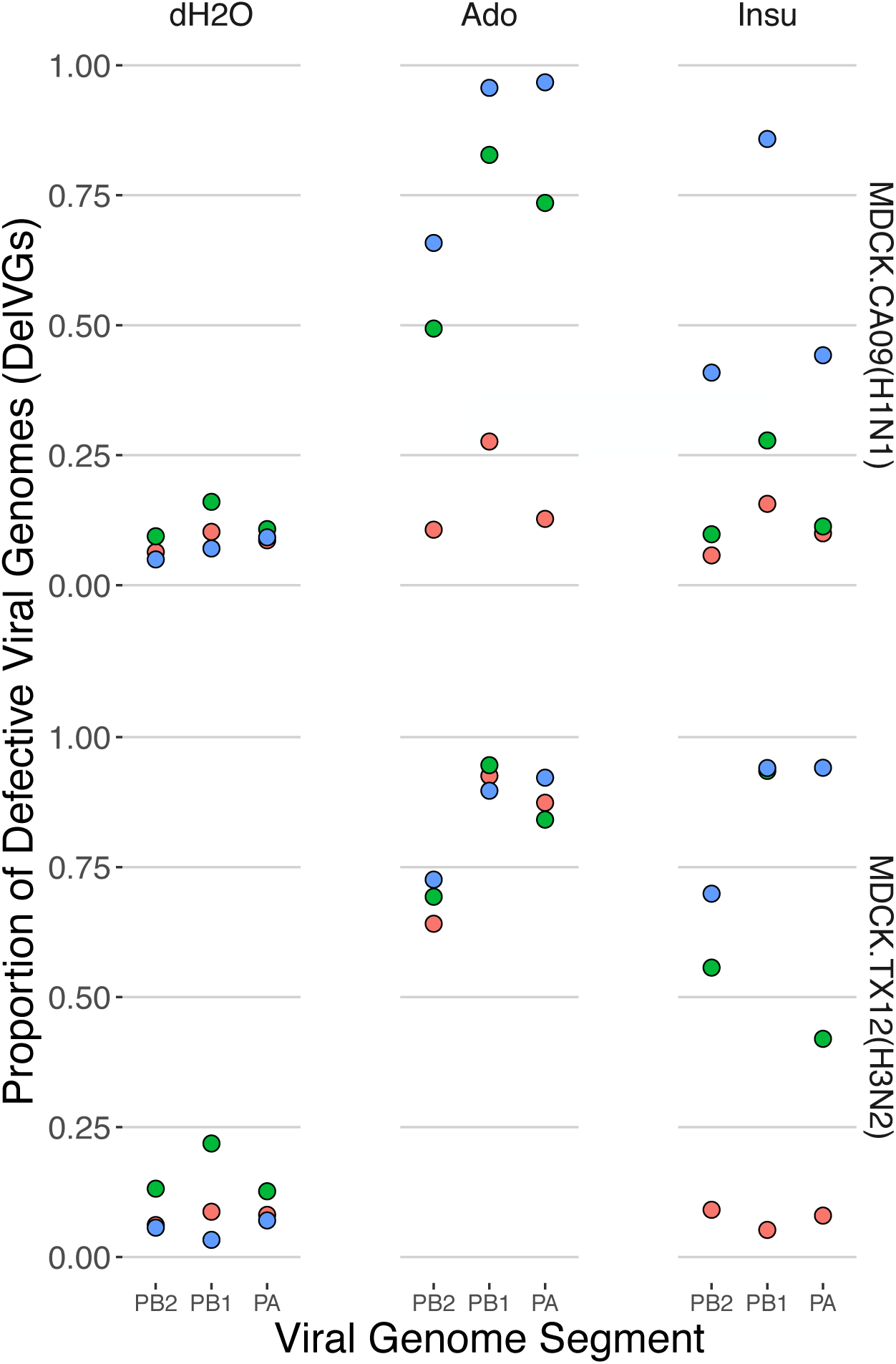
Adenosine and insulin increase polymerase complex defective viral genome segments. Production of influenza defective viral genomes (here measured as proportion of **Del**etion-containing **V**iral **G**enomes (DelVGs) after 18 hr infection; each point is the proportion of DelVGs in each influenza segment. Point colors represent data from three independent infections conducted on different days (i.e. bioreplicates).

### Insulin increases polymerase complex defective viral genomes

Insulin also increased the polymerase complex DelVG percentage, albeit with less consistency among strains than adenosine. For TX12, DelVGs of all three polymerase complex segments increased from 36.56% - 52.96% in a statistically significant manner. In CA09, DelVGs of all three polymerase complex segments also increased (**Figure 3**, **Table S1**), but these increases were not statistically significant.

### TCA cycle flux inhibitors 4-OI and UK5099 are potent amplifiers of total viral genomes across subtypes

The aerobic glycolysis inducers 4-OI and UK5099 increased overall (i.e. across all segments) TVG production in CA09 infections, as noted above (**Figure 4**, **Table S1**). Moreover, 4-OI increased DelVGs in seven out of eight segments in both strains in a statistically significant manner, with increases of 2,287 up to 11,798 total genomes compared to the DMSO control infections. In the remaining viral genome segment, NS, a statistically significant increase of 33,209 genomes was noted in CA09 infections. UK5099 also yielded statistically significant increases of 5,052 - 20,742 total viral genomes in both strains, but these increases were limited to four segments: HA, NP, NA, and M.

**Figure 4.**
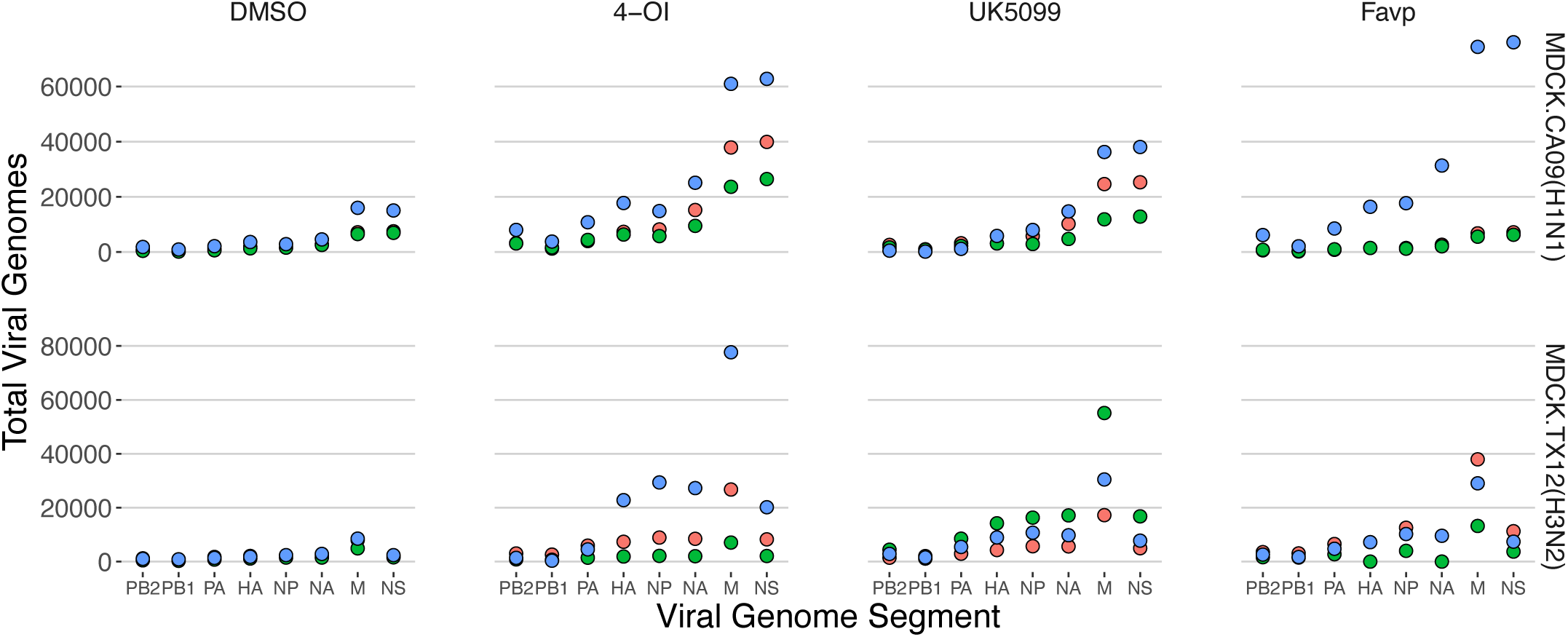
TCA cycle flux inhibitors 4-OI and UK5099 & purine analog favipiravir increase total viral genomes in A/H1N1 and A/H3N2 strains. Production of influenza **T**otal **V**iral **G**enomes (TVG) after 18 hr infection; each point is the count of total viral genomes (all segments included in each treatment). Point colors represent data from three independent infections conducted on different days (i.e. bioreplicates).

### The purine analog favipiravir increased total viral genomes across subtypes

Favipiravir infections showed a statistically significant average increase of 2,768 - 19,298 total viral genomes in segments PA, NP, and M in both CA09 and TX12 strains (**Figure 4**, **Table S1**). Favipiravir also showed a small, but statistically significant, 1.22% decrease in the percentage of DelVGs in TX12 infections (**Table S1**).

### Cyanobacterial extracts affected defective and total viral genomes in TX12 antigenic segments

TX12 infections responded to *Tolypothrix* extracts with a statistically significant increase of an average of 1,410 PB1 total viral genomes, whereas HA and NA registered statistically significant decreases of 3,178- 4,163 total viral genomes (**Figure 5**, **Table S1**). Additionally, *Tolypothrix* extracts reduced the NA DelVGs by a statistically significant albeit small 1.15% (**Table S1**). TX12 infections also had a curious response to *Leptolyngbya* extracts. There were substantial, statistically significant, *decreases* of 4,135-5,725 total HA and NA viral genome segments. These decreases led to near-zero recovery of HA and NA segments; less than 10-count of each segment in one bioreplicate, and zero in the remaining two bioreplicates (**Figure 5** blue box, **Table S1**). This effect was not an artefact experimentation; it occurred in three separate bioreplicates (run on different days) and supernatants of all treatments were processed individually—from RNA extraction, through successive RT-PCR and barcoding PCR amplification—before being pooled for sequencing library preparation.

**Figure 5.**
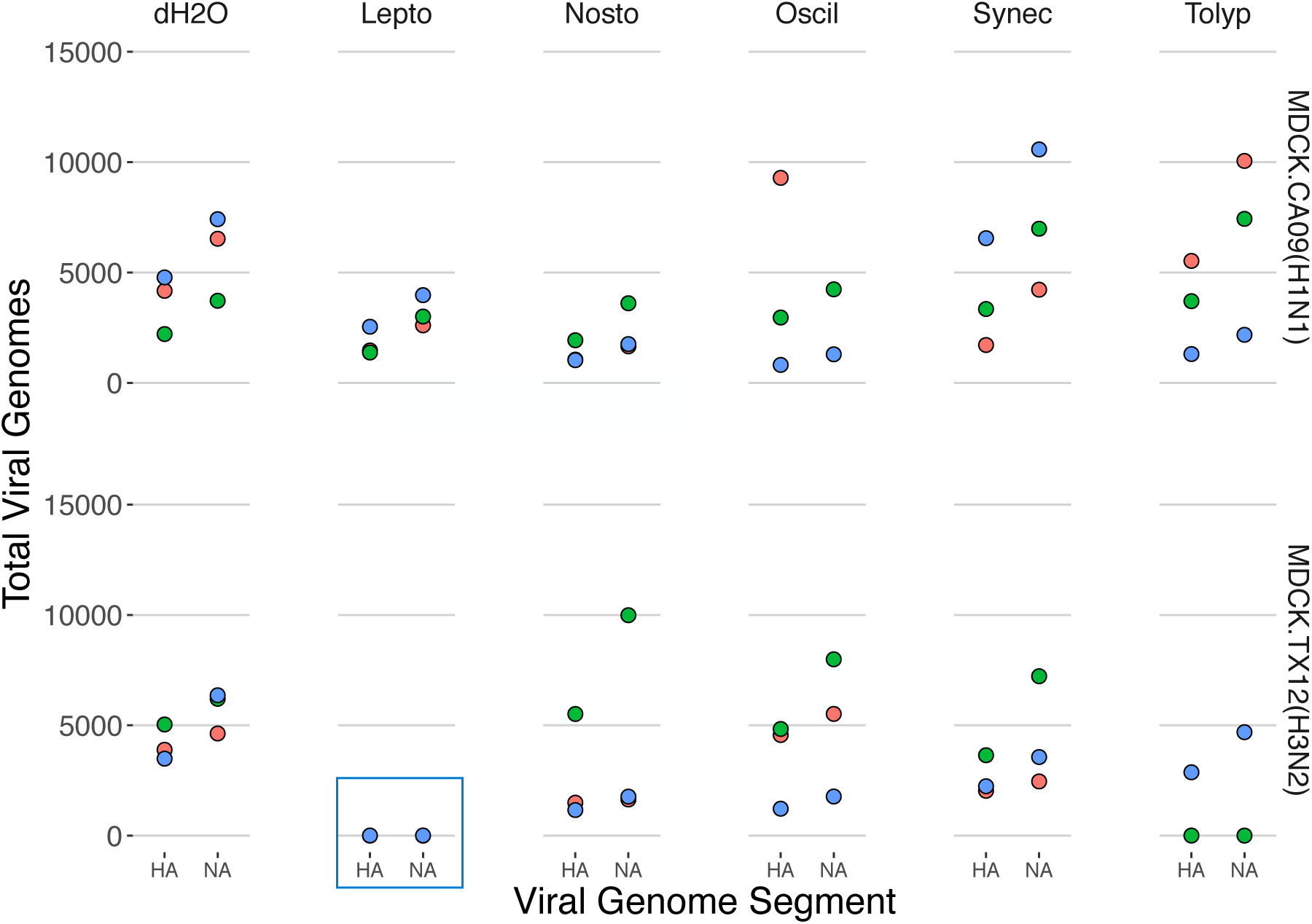
Cyanobacterial extracts suppress total viral genome production in A/H3N2 strain antigenic segments. Production of influenza **T**otal **V**iral **G**enomes (TVG) after 18 hr infection; each point is the count of total viral genomes in each influenza segment. Point colors represent data from three independent infections conducted on different days (i.e. bioreplicates).

## Discussion

Our study provides significant insights into the role of host metabolism in the production of defective viral genomes (DVGs) during Influenza A virus infection. The identification of adenosine, insulin, TCA cycle inhibitors, favipiravir, and cyanobacterial extracts as modulators of DVG production marks a crucial step toward understanding the de novo mechanisms underlying DVG production and exploring novel antiviral strategies.

### Adenosine is a potent amplifier of defective viral genome production

Adenosine emerged as a potent amplifier of DVG production across H1N1 and H3N2 subtypes. The significant increase in polymerase complex DelVGs in both strains suggests that adenosine has a broad-spectrum impact on DVG production. The notable increase in HA DelVGs in CA09 and NP DelVGs in TX12 highlights adenosine’s potential to differentially enhance DVG production across various genomic segments. Understanding the adenosine signaling cascade can shed light on the potential pathways of DVG production. For instance, the binding of extracellular adenosine to A1 G-protein coupled receptors on the MDCK cell plasma membrane triggers a phospholipase C (PLC)-dependent rise in cytosolic concentrations of inositol 1,4,5-triphosphate (IP3) and Ca^2+^, followed by a three-fold increase in the extracellular efflux of ATP which is dependent on the rapid sequestration of previously cytosolic Ca^2+^ into mitochondria (Migita, 2005; Migita, 2007). While extracellular ATP continues the signaling cascade in an autocrine and paracrine manner, Ca^2+^-sequestering mitochondria in contact with the endoplasmic reticulum (ER) engage in active trade of ions and membrane lipids (Hirabayashi, 2017). This interaction likely alters the ER lipid makeup and downstream vesicular trafficking to the plasma membrane, altering membrane rigidity and potentially driving the differential sorting of standard or defective genomes into progeny viral capsids (Blough, 1969; Blough, 1970; Lenard, 1976). Further research into these pathways will systematically reveal the host factors that reliably modulate the adenosine-associated increase in DelVG production.

### Insulin has strain-specific effects on polymerase complex DelVGs

Insulin demonstrated the ability to increase polymerase complex DelVGs, particularly in the TX12 strain. The statistically significant increases observed in TX12, contrasted with the non-significant increases in CA09, suggest a potential subtype-specific mechanism by which insulin signaling modulates DelVG production. Insulin binding to its receptor initiates a signaling cascade through multiple cellular pathways, including phosphoinositide 3-kinase (PI3K)-AKT and Ras-MAPK, which ultimately upregulates glucose uptake and glycolysis to fuel cell-wide anabolism in a HIF-1a and Myc-dependent manner (Hopkins, 2020). Further investigation into the molecular basis of this specificity could uncover critical insights into how host metabolic signals differentially influence viral genome dynamics across influenza subtypes and within segments of a given subtype.

### TCA Cycle inhibitors enhance total viral genome production

The TCA cycle inhibitors 4-OI and UK5099 significantly increased TVG production across multiple segments in both strains, highlighting a strong link between TCA cycle flux and viral genome replication. Specifically, the interference of normoxic pyruvate oxidation to CO2 gas by 4-OI, which inhibits mitochondrial succinate dehydrogenase (Daniels, 2019), and UK5099, which inhibits mitochondrial pyruvate carriers (Hildyard, 2005), led to significantly increased TVG production. This phenomenon is similar to the Warburg effect observed in tumor cells, where reduced-carbon biomass (pyruvate) is preserved to fuel proliferation (Lunt, 2011; Liu, 2019); in this case, the proliferation of viral genomes. While influenza virus is already known to induce an aerobic glycolysis state during normal pathogenesis (Ren, 2019; Ren, 2021), the addition of 4-OI or UK5099 appears to further enhance this metabolic state. Further research is needed to determine if 4-OI or UK5099 can also affect defective viral genome (DVG) production at different timepoints to fully explore their potential as broad-spectrum DVG-inducing drugs.

### Favipiravir increases total viral genome production

Favipiravir significantly increased TVGs across subtypes while slightly reducing DelVG proportions in TX12 infections. Considering favipiravir’s known role as a mutagen and premature terminator of RNA elongation during influenza replication, the observed increase in total genomes suggests that our chosen dose of favipiravir was insufficient to induce lethal mutagenesis. This hypothesis is supported by two key points: (i) the 1 µM dose of favipiravir used in our study is an order of magnitude lower than its 11 µM EC50 concentration (Vanderlinden, 2016), and (ii) 1 µM favipiravir is easily outcompeted by the 100-200 µM basal intracellular concentration of its nucleotide competitor GTP (Zala, 2017). The observed permissive mutagenesis by favipiravir, coupled with increased TVGs and slightly reduced DVGs, suggests that misincorporation-based mutation may not be a necessary precursor to DelVG production. Further research is needed to better understand the relationship between nucleotide misincorporation and DVG production.

### Cyanobacterial extracts suppress production of antigenic segments in tx12 (a/h3n2)

The differential response of TX12 segments to cyanobacterial extracts, particularly the substantial decreases in HA and NA segments with *Leptolyngbya* and *Tolypothrix* extracts, highlights the potential of natural products to modulate viral genome dynamics. This underscores the need for further exploration into the active components of these extracts and their mechanisms of action (Singh, 2017; Silva, 2018; Mazur-Marzec, 2021; Tiwari, 2020; Ferrazano, 2020). Notably, the near absence of HA and NA segments in response to *Leptolyngbya* extracts suggests their active exclusion from entry into progeny capsids. This finding points to a non-random, segment-selective force acting on viral segments during replication, cytoskeletal trafficking, and/or progeny capsid assembly, warranting further investigation into the underlying mechanisms.

### Implications for future research and therapeutic development

Overall, our findings reinforce the critical role of host metabolism in influencing DVG production and viral pathogenesis. Our findings suggest that the social interactions observed between defective and full-length viral genomes, can be crucially influenced by the host context (Leeks et al. 2023; Díaz-Muñoz et al., 2017). The identification of metabolic effectors that modulate DVG and TVG production provides a foundation for future research aimed at uncovering the molecular mechanisms driving these effects. Understanding the shared characteristics of these inducers will be pivotal in developing highly targeted antiviral therapies that exploit metabolic vulnerabilities of the influenza virus. Additionally, understanding the role of host metabolism in DelVG production could help to predict how various host states in flu patients (such as diabetes, cancer, pregnancy or other metabolic states) can shape viral evolution and influence disease severity (Ghedin, 2017). For instance, pregnancy, known to lead to severe flu infections, also leads to the selection of specific mutations that increase pathogencity in H1N1 strains (Engels, 2017). Thus, understanding how metabolic states pre-dispose flu infections to produce more or less DelVGs or TVGs, which can influence disease severity, could be an important clinical tool. In conclusion, this study advances our understanding of the interplay between host metabolism and Influenza A virus genome dynamics. By identifying key metabolic modulators of DVG production, we pave the way for innovative therapeutic strategies that leverage defective interference to combat influenza infections. Future investigations should focus on the detailed molecular pathways involved and the potential for these metabolic effectors to be integrated into broad-spectrum antiviral treatments.

## Acknowledgements

This project was funded by NIH grants 4R00AI119401-02 and 1R01AI179873-01 to SDM. IA was supported by NIH grant 4R00AI119401-02 and 1R01AI179873-01 to SDM and a Craft Consult Biotechnology Dissertation Fellowship. John Albeck and Enoch Baldwin provided helpful feedback on this manuscript. Ted Ross kindly provided Influenza A strains from his collection. John Meeks graciously provided cyanobacteria from his collection.

