## Supplementary Materials for "Influenza A defective viral genome production is altered by metabolites, metabolic signaling molecules, and cyanobacteria extracts"

### Extended Results

#### *Drugs affect DVG proportion and TVG production at the segment level for CA09 and TX12*

Continued from Results in the main text, we report the segment-specific effects of drug treatments on DelVG proportion and total genomes that were significant relative to their vehicle control (DMSO, $dH_2O$). As stated, we analyzed differences among treatments using ANOVA, adding bioreplicate, strain, and segment as covariates.

| Strain | Vehicle | Measure | Segment | Treatment | Estimate | Std. Error | t value | Pr(>\|t\|) | |
|---|---|---|---|---|---|---|---|---|---|
| **CA09** | H2O | PropDVG | PB1 | TxAdo | 0.575611 | 0.204348 | 2.817 | 1.24E-02 | * |
| **CA09** | H2O | PropDVG | PB2 | TxAdo | 0.350356 | 0.10737 | 3.263 | 0.00488 | ** |
| **CA09** | H2O | PropDVG | PA | TxAdo | 0.514519 | 0.140622 | 3.659 | 0.00212 | ** |
| **TX12** | H2O | PropDVG | PB1 | TxAdo | 0.80981 | 0.20321 | 3.985 | 1.06E-03 | * |
| **TX12** | H2O | PropDVG | PB2 | TxAdo | 0.603432 | 0.122571 | 4.923 | 0.000153 | ** |
| **TX12** | H2O | PropDVG | PA | TxAdo | 0.78621 | 0.178854 | 4.396 | 0.000451 | *** |
| **TX12** | H2O | PropDVG | NP | TxAdo | 0.009966 | 0.0036557 | 2.726 | 0.014943 | * |
| **CA09** | H2O | PropDVG | HA | TxAdo | 0.337257 | 0.1237154 | 2.726 | 0.015 | * |
| **TX12** | H2O | PropDVG | PB1 | TxInsu | 0.52964 | 0.20321 | 2.606 | 0.01909 | * |
| **TX12** | H2O | PropDVG | PA | TxInsu | 0.387404 | 0.178854 | 2.166 | 0.045756 | * |
| **TX12** | H2O | PropDVG | PB2 | TxInsu | 0.365633 | 0.122571 | 2.983 | 0.008785 | ** |
| **TX12** | H2O | PropDVG | NP | TxInsu | 0.007798 | 0.0036557 | 2.133 | 0.048746 | * |
| **TX12** | DMSO | PropDVG | NA | TxFavp | -0.01216 | 0.0047605 | -2.556 | 0.0286 | * |
| **TX12** | H2O | PropDVG | NA | TxLepto | -0.01946 | 0.004713 | -4.13 | 0.001021 | ** |
| **TX12** | DMSO | PropDVG | NA | TxMK2206 | 0.014262 | 0.0047605 | 2.996 | 0.0134 | * |
| **TX12** | H2O | PropDVG | NA | TxTolyp | -0.01150 | 0.004713 | -2.441 | 0.028536 | * |
| **BOTH** | DMSO | TVG | PB2 | Tx4-OI | 2287.3 | 905.6 | 2.526 | 0.0177 | * |
| **BOTH** | DMSO | TVG | PB1 | Tx4-OI | 1177.67 | 462.2 | 2.548 | 0.0168 | * |
| **BOTH** | DMSO | TVG | PA | Tx4-OI | 3886.3 | 1285.5 | 3.023 | 0.00543 | ** |
| **BOTH** | DMSO | TVG | HA | Tx4-OI | 8590 | 2447.24 | 3.51 | 0.00159 | ** |
| **BOTH** | DMSO | TVG | NP | Tx4-OI | 9482.83 | 2792.07 | 3.396 | 0.00213 | ** |
| **BOTH** | DMSO | TVG | NA | Tx4-OI | 11798.3 | 3463.2 | 3.407 | 0.00207 | ** |
| **BOTH** | DMSO | TVG | M | Tx4-OI | 30476 | 8857 | 3.441 | 0.0019 | ** |
| **CA09** | DMSO | TVG | NS | Tx4-OI | 33209 | 12276 | 2.705 | 0.0221 | * |
| **BOTH** | DMSO | TVG | HA | TxUK5099 | 5051.67 | 2447.24 | 2.064 | 0.04873 | * |
| **BOTH** | DMSO | TVG | NP | TxUK5099 | 6189.17 | 2792.07 | 2.217 | 0.03526 | * |
| **BOTH** | DMSO | TVG | NA | TxUK5099 | 7577.3 | 3463.2 | 2.188 | 0.0375 | * |
| **BOTH** | DMSO | TVG | M | TxUK5099 | 20742 | 8857 | 2.342 | 0.0268 | * |
| **BOTH** | DMSO | TVG | PA | TxFavp | 2767.7 | 1285.5 | 2.153 | 0.04041 | * |
| **BOTH** | DMSO | TVG | NP | TxFavp | 5809.5 | 2792.07 | 2.081 | 0.04708 | * |
| **BOTH** | DMSO | TVG | M | TxFavp | 19298 | 8857 | 2.179 | 0.0382 | * |

| | | | | | | | | | |
|---|---|---|---|---|---|---|---|---|---|
| **TX12** | H2O | TVG | HA | TxLepto | -4135.67 | 1052.26 | -3.93 | 0.001336 | ** |
| **TX12** | H2O | TVG | NA | TxLepto | -5725.33 | 1824.47 | -3.138 | 0.00726 | ** |
| **TX12** | H2O | TVG | PB1 | TxTolyp | 1410 | 532.93 | 2.646 | 0.0176 | * |
| **TX12** | H2O | TVG | HA | TxTolyp | -3178 | 1052.26 | -3.02 | 0.008611 | * |
| **TX12** | H2O | TVG | NA | TxTolyp | -4163.67 | 1824.47 | -2.282 | 0.03864 | * |

**Supplemental Table 1**. **Drugs significantly alter total viral genomes and proportion of deletion containing viral genomes at the segment level**. Segment-specific statistically significant predictors of the proportion of **Del**etion-containing **V**iral **G**enomes (DelVGs) and **T**otal **V**iral **G**enomes (TVG) and their parameter estimates from ANOVA.

### *Drugs affect DVG proportion and TVG Production at the genome level for CA09 and TX12*

Below are visualizations of DVG relative abundance (**Supplementary Figure 1**) and total viral genomes (**Supplementary Figure 2**) for CA09 and TX12 *at the genome level*, averaged across three bioreplicates.

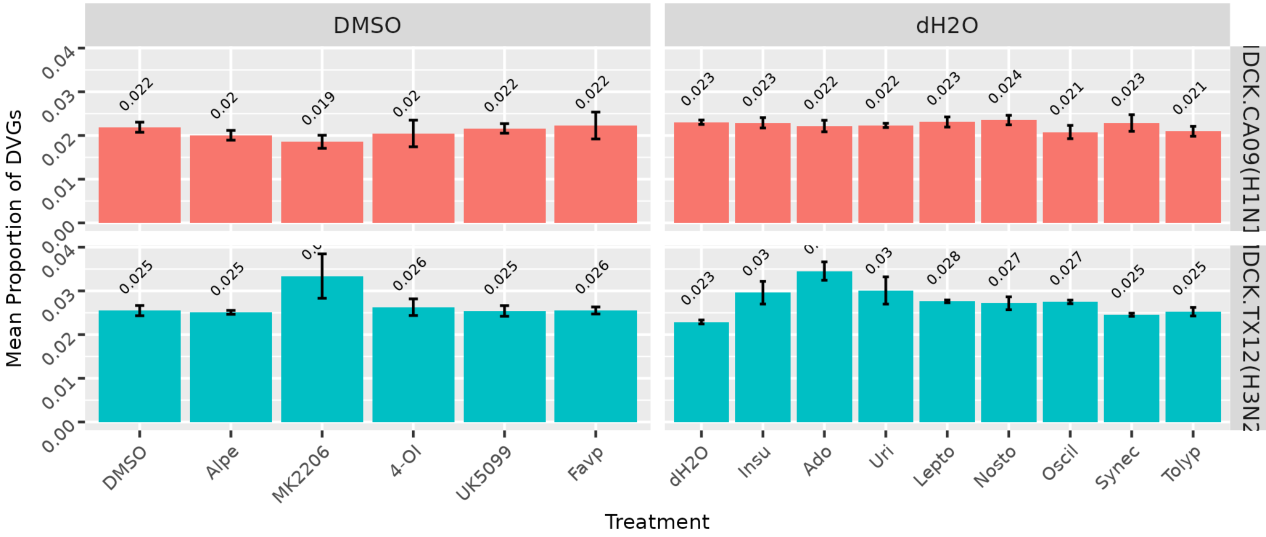


**Supplemental Figure 1. Mean proportion of total viral genomes (CA09/TX12) that are DVGs after 18 h.p.i. under different treatment conditions**; no trypsin. Vehicle treatment groups received either DMSO or $dH_2O$ treatment. n = 3 bioreplicates, sem.

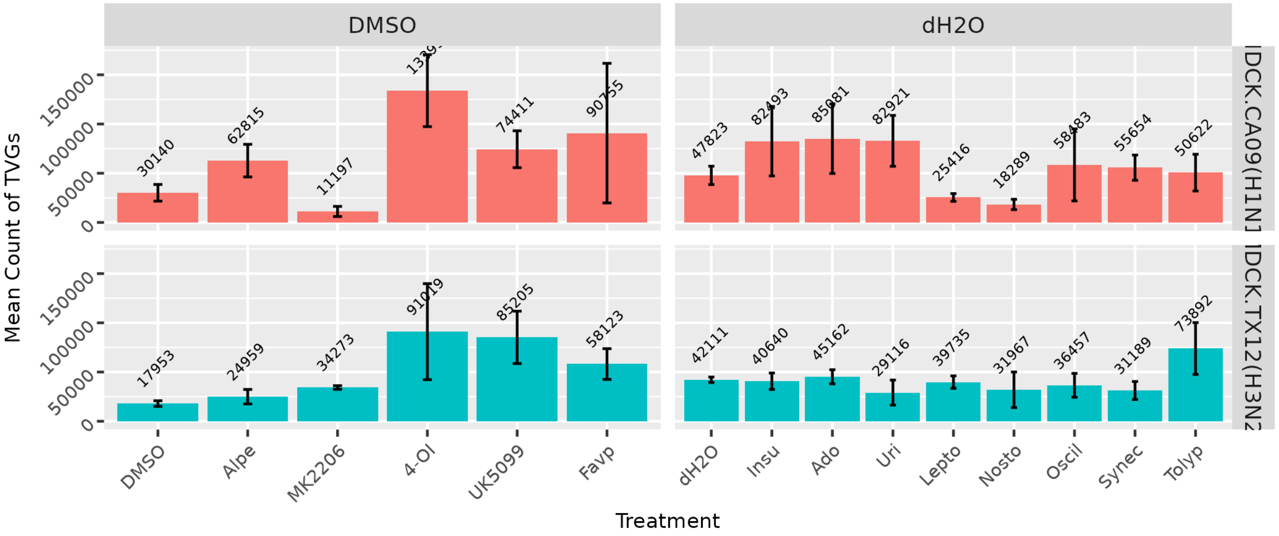


**Supplemental Figure 2. Mean count of total viral genomes (CA09/TX12) recovered at 18 h.p.i. under different treatment conditions**; no trypsin. Vehicle treatment groups received either DMSO or $dH_2O$ treatment. n = 3 bioreplicates, sem.

**3A**

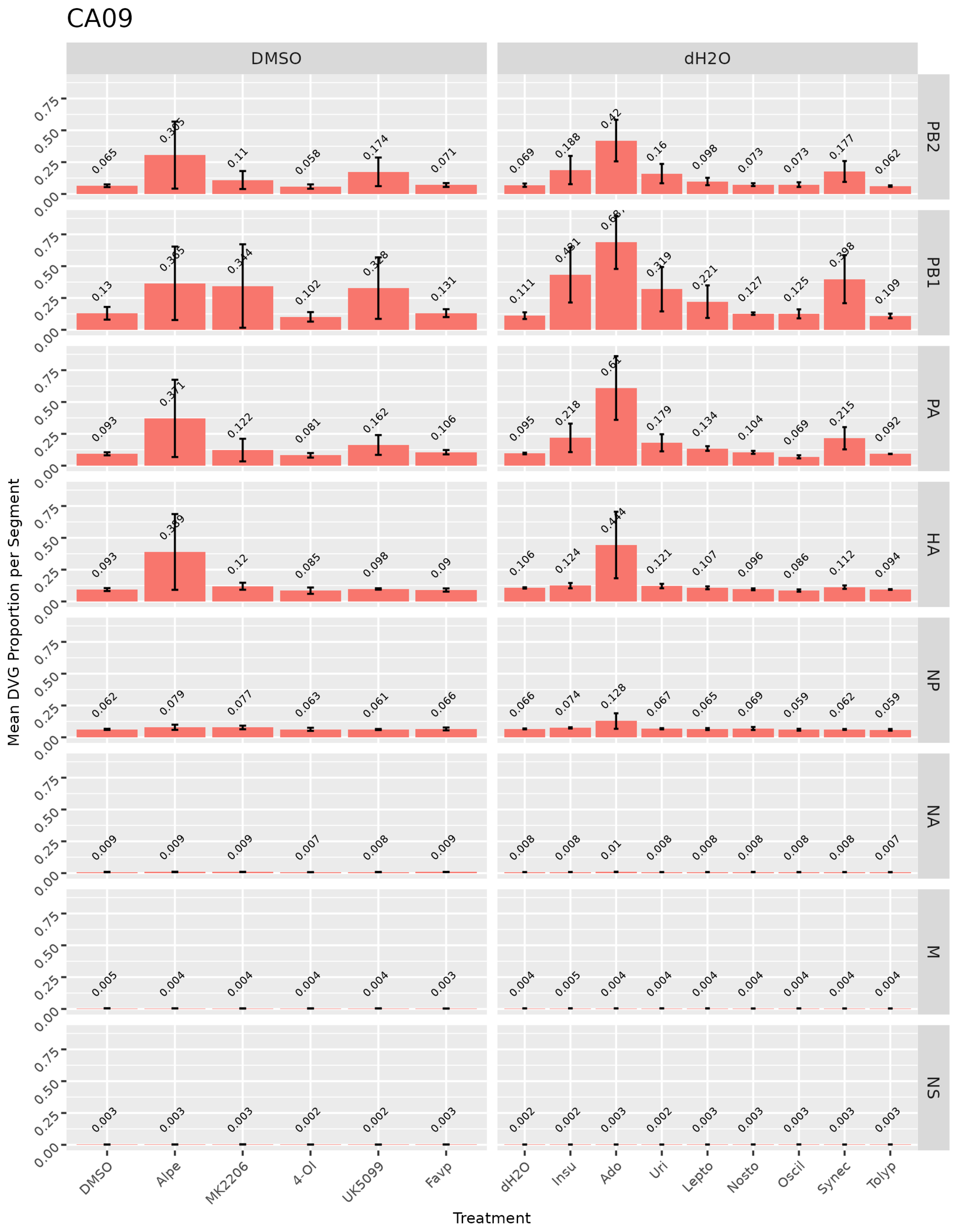

CA09
DMSO
dH2O
PB2
PB1
PA
HA
NP
NA
M
NS
Mean DVG Proportion per Segment
Treatment
DMSO
Alpe
MK2206
4-OI
UK5099
Favp
dH2O
Insu
Ado
Uri
Lepto
Nosto
Oscil
Synec
Tolyp
0.00
0.25
0.50
0.75

**3B**

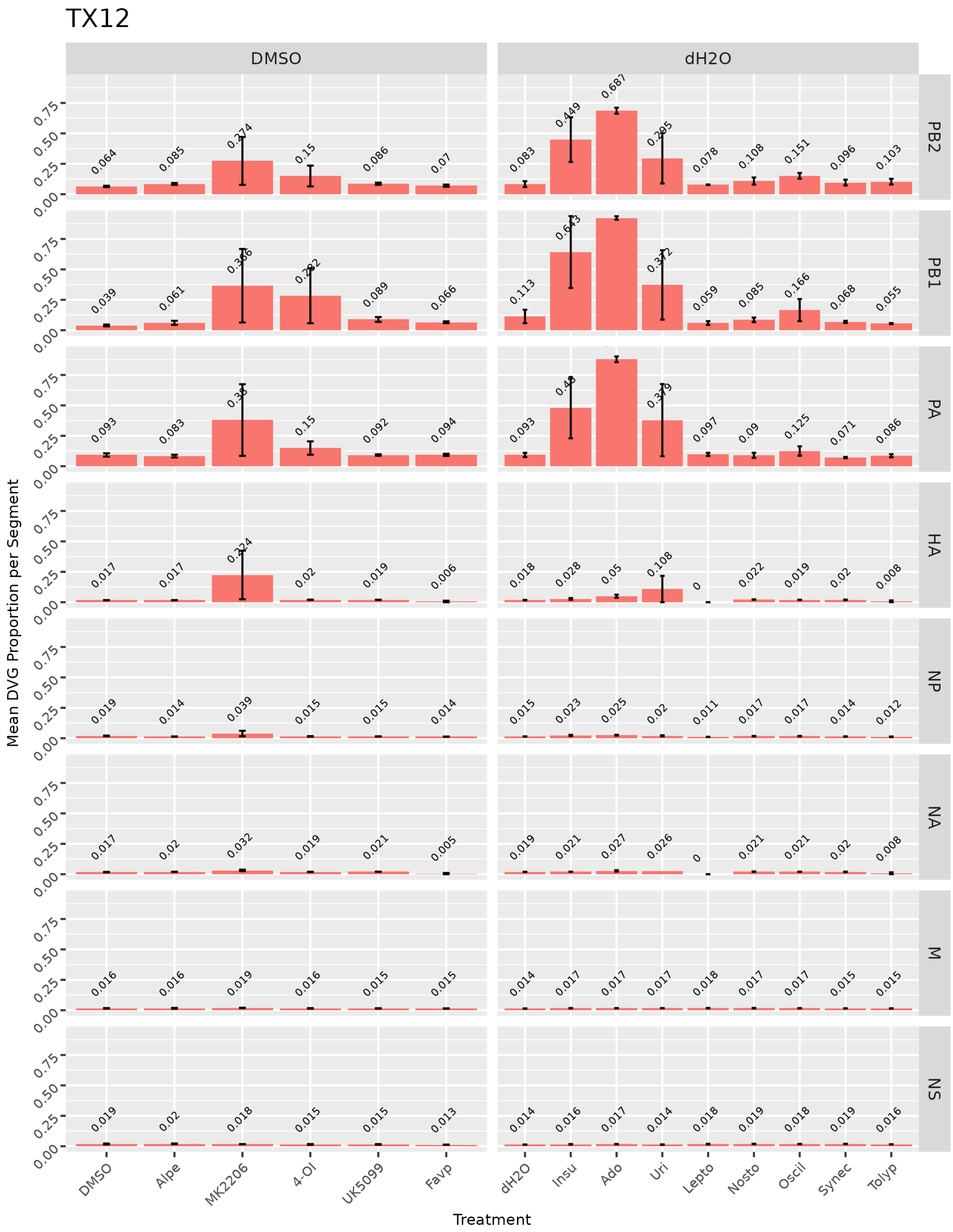


**Supplementary Figure 3. Mean proportion of total viral genomes per segment that are DVGs at 18 h.p.i. under different treatment conditions**; no trypsin. (**A**) CA09. (**B**) TX12. Vehicle treatment groups received either DMSO or $dH_2O$ treatment. n = 3 bioreplicates, sem.

**3.4A**

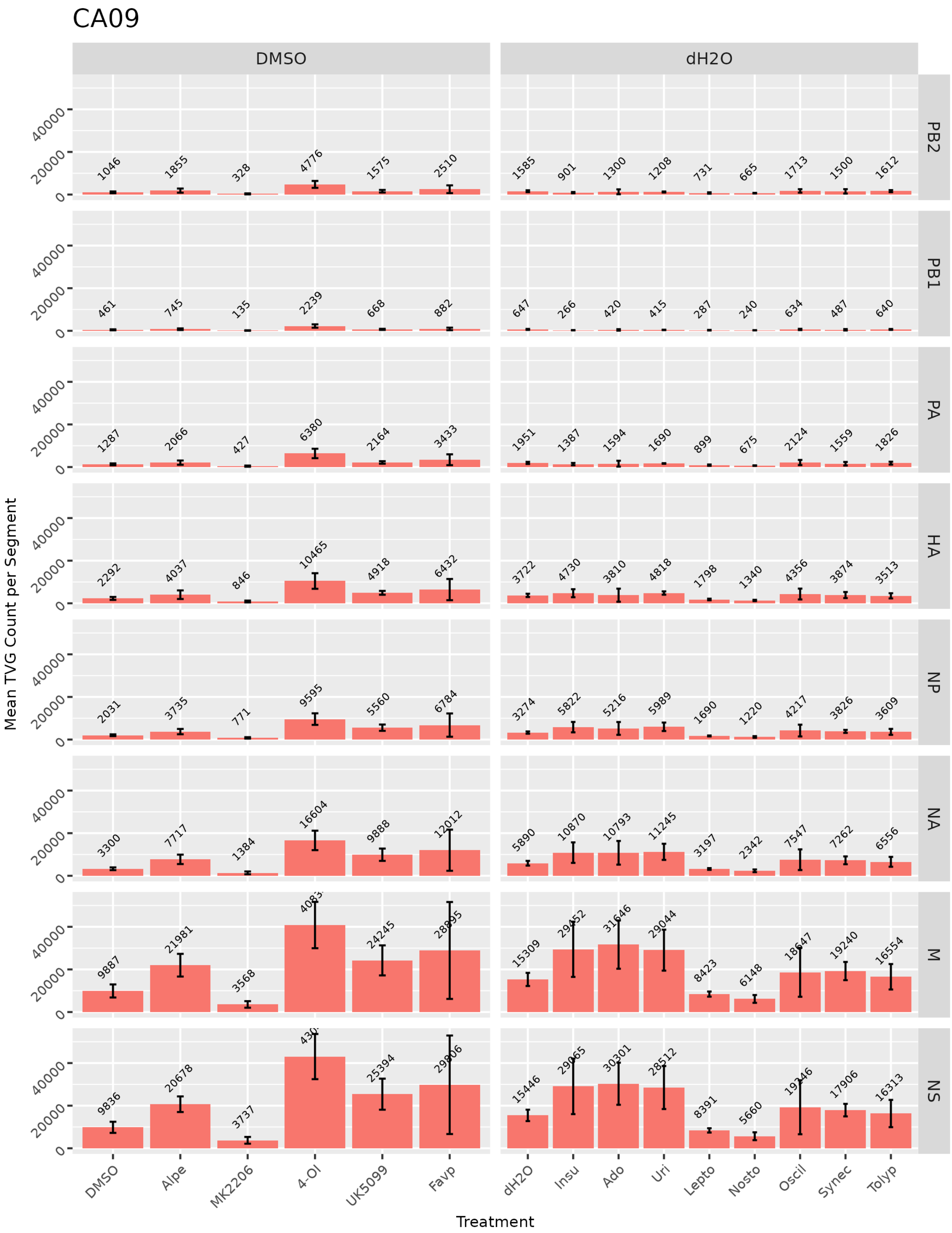

CA09
DMSO
dH2O
PB2
PB1
PA
HA
NP
NA
M
NS
Mean TVG Count per Segment
Treatment
DMSO
Alpe
MK2206
4-OI
UK5099
Favp
dH2O
Insu
Ado
Uri
Lepto
Nosto
Oscil
Synec
Tolyp

**3.4B**

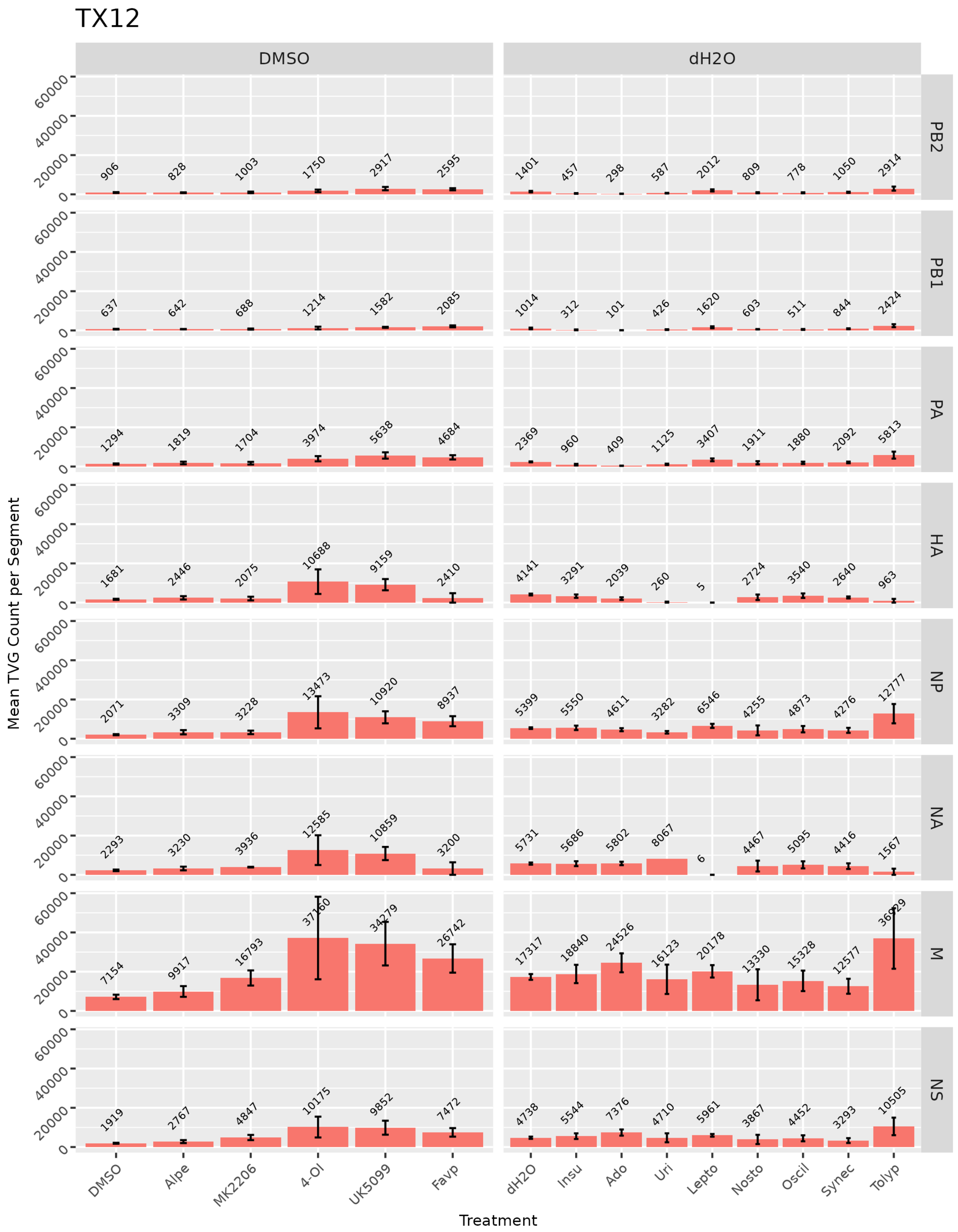


**Supplementary Figure 4. Mean count of total viral genomes per segment produced after 18 h.p.i. under different treatment conditions**; no trypsin. (**A**) CA09. (**B**) TX12. Vehicle treatment groups received either DMSO or $dH_2O$ treatment. n = 3 bioreplicates, sem.
